# Essential role for an isoform of *Escherichia coli* translation initiation factor IF2 in repair of two-ended DNA double-strand breaks

**DOI:** 10.1101/2021.11.18.469101

**Authors:** Jillella Mallikarjun, J Gowrishankar

## Abstract

In *Escherichia coli*, three isoforms of the essential translation initiation factor IF2 (IF2-1, IF2-2, and IF2-3) are generated from separate in-frame initiation codons in *infB*. The isoforms have earlier been suggested to additionally participate in DNA damage repair and replication restart. It is also known that the proteins RecA and RecBCD are needed for repair of DNA double-strand breaks (DSBs) in E. *coli*. Here we show that strains lacking IF2-1 are profoundly sensitive to two-ended DSBs in DNA generated by radiomimetic agents phleomycin or bleomycin, or by endonuclease I-SceI. However, these strains remained tolerant to other DSB-generating genotoxic agents or perturbations to which *recA* and *recBC* mutants remained sensitive, such as to mitomycin C, type-2 DNA topoisomerase inhibitors, or DSB caused by palindrome cleavage behind a replication fork. Data from genome-wide copy number analyses following I-SceI cleavage at a single chromosomal locus suggested that, in a strain lacking IF2-1, the magnitude of break induced replication through replication restart mechanisms is largely preserved but the extent of DNA resection around the DSB site is reduced. We propose that in absence of IF2-1 it is the annealing of a RecA nucleoprotein filament to its homologous target that is weakened, which in turn leads to a specific failure in assembly of *Ter-to-oriC* directed replisomes needed for consummation of two-ended DSB repair.

**Importance:** Double-strand breaks (DSBs) in DNA are major threats to genome integrity. In *Escherichia coli*, DSBs are repaired by Rec- and RecBCD-mediated homologous recombination (HR). This study demonstrates a critical role for an isoform (IF2-1) of the translation initiation factor IF2 in the repair of two-ended DSBs in *E. coli* (that can be generated by ionizing radiation, certain DNA-damaging chemicals, or endonuclease action). It is proposed that IF2-1 acts to facilitate the function of RecA in the annealing between a pair of DNA molecules during HR.

## Introduction

Amongst the most severe forms of damage to the genetic material in all life forms is the double strand break (DSB) in DNA (1–11). In bacteria such as *Escherichia coli*, repair of a DSB is achieved by homologous recombination (HR), which entails a step of synapsis between the broken DNA ends and an intact DNA duplex (1, 2, 7, 10, 11). RecA is the central protein of HR that mediates synapsis, and its action in DSB repair is preceded by a set of (pre-synaptic) reactions performed by the RecBCD complex.

RecBCD is a dual function exonuclease-helicase that acts on broken DNA ends to generate 3’-overhangs of single-stranded (ss) DNA, upon which the RecA nucleoprotein filaments are then assembled. Formation of these 3’-overhangs is facilitated by the presence of *cis* sites in DNA designated Chi (which is an asymmetric 8-bp sequence), since the exonuclease (but not helicase) activity of RecBCD is attenuated upon its encounter with a Chi site (1–3, 10, 12).

The RecA-mediated synaptic step in DSB repair results in formation of D-loops and Holliday junctions that serve as substrates for action by the RuvABC proteins (1, 2, 13). This is followed by a process of replication restart, by which a replisome is assembled on the post-synaptic structures to enable integration of the broken DNA ends into the circular bacterial chromosome. The proteins PriABC, DnaT, and Rep are proposed to act through multiple alternative and redundant pathways to achieve replication restart (13–18).

As may be expected from the description above, deletion mutations in genes *recA, recB, recC, ruv* or *priA* confer marked sensitivity to DSBs in DNA (1–3). On account of redundancy in the replication restart pathways, mutants individually deficient for PriB, PriC, or the helicase activity of PriA (through a *priA300* mutation) are tolerant to DSBs, but the double mutants *priB-priC* or *priB-priA300* are sensitive (13–18).

DSBs in *E. coli* may be either one-ended or two-ended (2, 13). The former occur at sites of replication fork collapse or disintegration, in which one sister chromatid arm is dissociated from the remaining circular chromosome (1, 2, 13, 19). RecA- and RecBCD-mediated repair of a one-ended DSB results in a restart replisome that is directed to progress toward the chromosomal terminus region (*oriC-*to-*Ter* direction).

Two-ended DSBs occur when there is a clean break of the DNA duplex unrelated to replication, as with ionizing radiation, radiomimetic agents (20, 21) such as phleomycin (Phleo) or bleomycin (Bleo), or cleavage by endonucleases such as restriction enzymes or I-SceI. The general concept has been that repair of two-ended DSBs in *E. coli* “can [simply] be viewed as a combination of two initially independent” events of one-ended DSB repair (1), by which a pair of restart replisomes are established that converge toward one another in *oriC-*to-*Ter* and *Ter-to-oriC* directions, respectively (“ends-in” replication) (2, 7, 13). In many other life forms, distinctive mechanisms apparently exist for two-ended DSB repair (22–26).

We now report that isoforms of the translation initiation factor IF2 can specifically modulate two-ended DSB repair in *E. coli*. These findings are complementary to those reported by us in the accompanying paper (27), that the IF2 isoforms can influence HR through RecA and the RecFORQ-mediated pre-synaptic pathway. IF2 is an essential protein for translation initiation (28), which exists in three isoforms (IF2-1, IF2-2, and IF2-3) that are generated from three in-frame initiation codons 1, 158, and 165 of the *infB* ORF (29–32). Any one of these isoforms is sufficient for the translation initiation function.

Nakai and coworkers had previously reported that DNA transactions including damage repair processes are impacted upon differential expression of IF2 isoforms in the cells, and they had suggested that this is a consequence of the isoforms’ effects on different pathways of replication restart (33–35). Based on our present studies involving genome-wide copy number analyses following a two-ended DSB at a single chromosomal locus, we propose an alternative model wherein absence of isoform IF2-1 abrogates the *Ter-*to-*oriC* replication component that is needed for two-ended DSB repair.

## Results

### Loss of IF2-1 confers profound sensitivity to two-ended DSBs in DNA

Given our findings in the accompanying paper (27) that IF2 isoforms can modulate HR functions, as well as on earlier reports from the Nakai group (34, 35) on differential sensitivity to DNA damage of strains deficient for particular IF2 isoforms, we investigated in more detail the relationship between expression of the IF2 isoforms and DNA damage tolerance.

The following nomenclature is employed below for different alleles and constructs related to IF2 isoforms, as was also adopted in the accompanying paper (27). *infB*^+^ and Δ*infB* refer to the wild-type and deletion alleles, respectively, at the native chromosomal location. For the set of three ectopic *infB* chromosomal constructs described by Nakai and coworkers (34, 35) (in each of which IF2 expression remains under control of the natural *cis* regulatory elements for *infB*), Δ*2*,*3* refers to that encoding only IF2-1, but not IF2-2 or IF2-3; Δ*1* to that encoding both IF2-2 and IF2-3, but not IF2-1; and Δ*Nil* to that encoding all three isoforms. For plasmid-encoded *infB* constructs under control of the isopropyl-β-D-thiogalactoside (IPTG)-induced P_*trc*_ promoter, P_*trc*_-Δ*1* refers to that expressing both IF2-2 and IF2-3, but not IF2-1; and P_*trc*_-Δ*1*,*2* to that expressing IF2-3 alone.

Our results indicate that the Δ*1* strain (deficient for IF2-1, and expressing just the IF2-2,3 isoforms) is profoundly sensitive to perturbations that unambiguously generate two-ended DSBs on the chromosome. These perturbations include exposure to radiomimetic agents Phleo or Bleo, and cleavage by endonuclease I-SceI. Thus, at concentrations of Phleo or Bleo that provoke chromosomal two-ended DSBs (and so render a *recA* mutant inviable), the strain lacking IF2-1 was killed to at least the same extent as *recA* itself; on the other hand, the Δ*2*,*3*, *priA300* or Δ*priB* strains were as tolerant to these agents as was the Δ*Nil* strain (Fig. 1A). Phleo sensitivity of the Δ*1* strain could be complemented by plasmid-borne *infB^+^* (Supp. Fig. S1A). It is the RecBCD pre-synaptic pathway that is involved in RecA-mediated recombinational repair of two-ended DSBs in DNA (1–5), which was confirmed also in our experiments by demonstrating that a *recB* but not *recO* mutant is sensitive to Phleo and Bleo (Supp. Fig. S1B).

**Figure 1:**
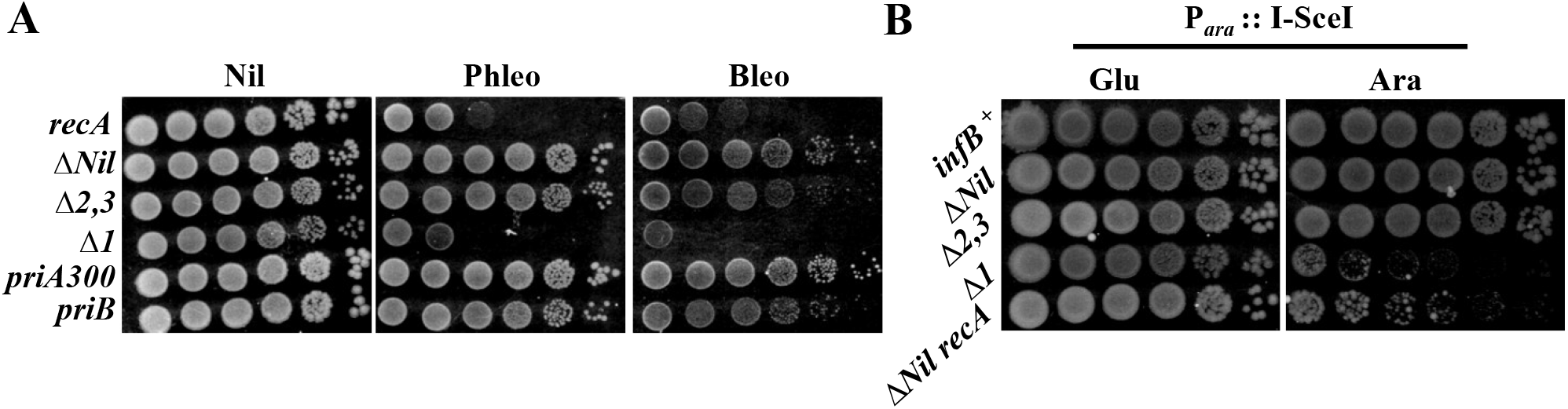
Loss of IF2-1 is associated with sensitivity to two-ended DSBs in DNA. In both panels, strains designated Δ*Nil*, Δ*1*, or Δ*2*,*3* were also Δ*infB*. Dilution-spotting assays were performed on LB medium with supplements as indicated on top (Phleo and Bleo each at 0.5 μg/ml, Glu and Ara each at 0.2%), of different strains whose relevant genotypes/features are marked. Strains employed for different rows were (from top, all strain numbers are prefixed with GJ): panel A – 19844, 19193, 19194, 15494, 15495, and 19812; and panel B – 15837, 19804, 19805, 19806, and 19818.

Likewise, the Δ*1* strain (but not Δ*Nil* or Δ*2*,*3*), as well as the *recA* and *recB* derivatives of Δ*Nil*, were markedly sensitive to two-ended DSB at an I-SceI site in the *lacZ* locus, that was generated by L-arabinose (Ara)-induced expression of endonuclease I-SceI (Fig. 1B; and Supp. Fig. S2 ii). These results therefore establish that loss of IF2-1 is associated with compromise of two-ended DSB repair mediated by RecA and the RecBCD pre-synaptic pathway. On the other hand, none of the *pri* mutations tested (*priA300*, Δ*priB* or Δ*priC*) had any effect on sensitivity or tolerance to I-SceI cleavage of the Δ*Nil*, Δ*1*, or Δ*2*,*3* strains (Supp. Fig. S2, compare i with iii-v).

Interestingly, with lower concentrations of Phleo or Bleo wherein *recA* viability was only marginally affected, the strain without IF2-1 continued to exhibit marked sensitivity (Supp. Fig. S1C, compare rows 4 and 7 of three panels at left; supported also by data in Bleo sub-panel of Fig. 1A). This would suggest that at the low doses, DNA damage other than DSBs continues to occur (perhaps ss-DNA gaps), which can be repaired by RecA-independent mechanism(s) [such as by DNA polymerase I followed by DNA ligase (1)] but only so if IF2-1 is present.

### Confirmation by flow cytometry of Δ*1*’s sensitivity to two-ended DSBs

Sensitivity of the Δ*1* derivatives to Phleo or to I-SceI cleavage was demonstrated also by flow cytometry following propidium iodide staining for dead cells in cultures, wherein these strains exhibited much greater cell death than did the isogenic Δ*Nil* or Δ2,3 strains (Fig. 2, middle and bottom rows). Even in ordinarily grown cultures without any DNA damaging agent or treatment, around 8% of cells of the Δ*1* strain were scored as dead, a value comparable to that for *recA* and much higher than those for Δ*Nil* or Δ*2*,*3* derivatives (< 0.5% each) (Fig. 2, top row); it should be noted, however, that these values do not take into account any contributions to inviability by anucleate cells or cell lysis in the cultures (36, 37).

**Figure 2:**
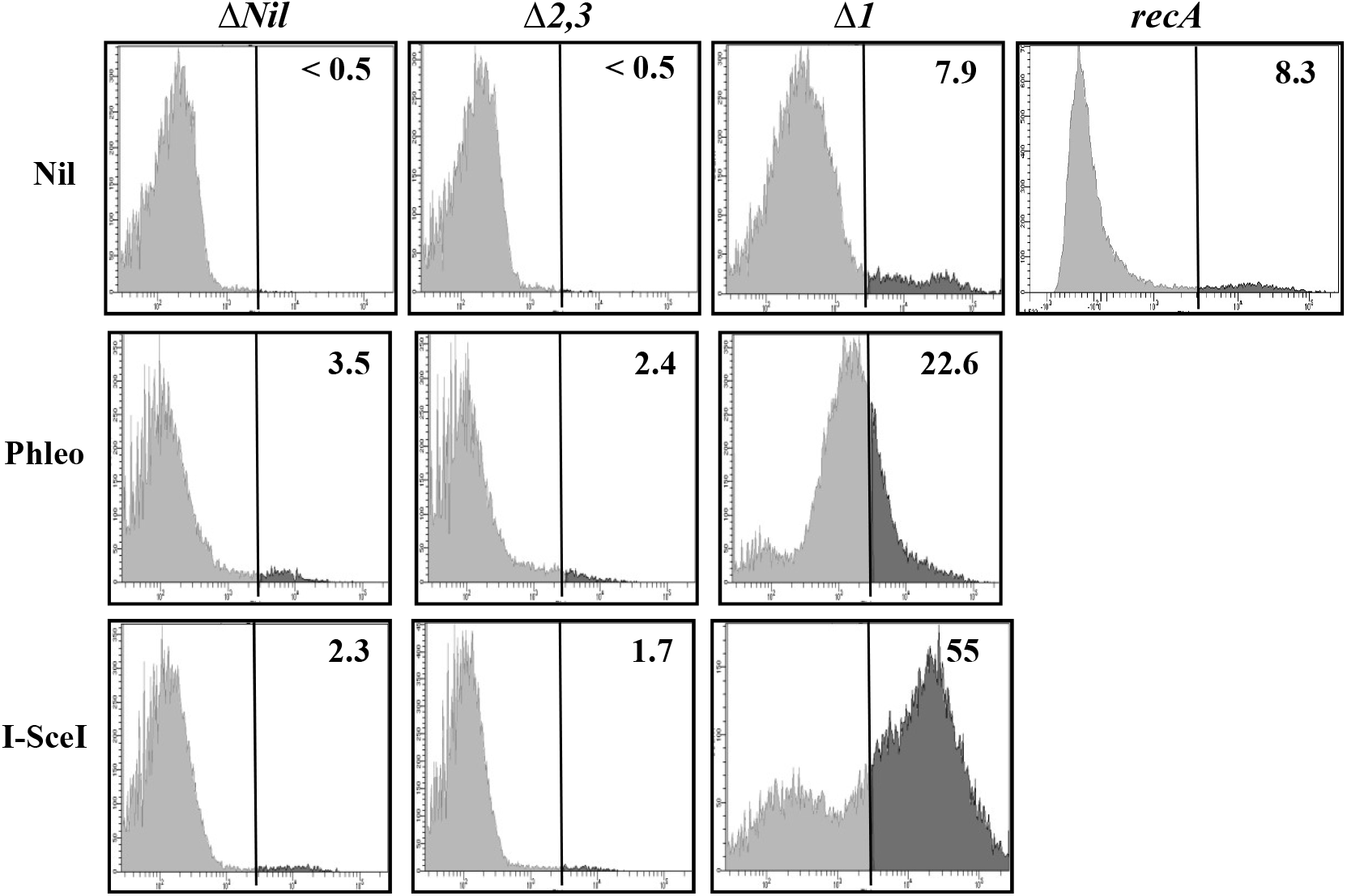
Demonstration by flow cytometry of sensitivity to two-ended DSBs in DNA in absence of IF2-1. Flow cytometry was performed following propidium iodide staining of cells in LB-grown cultures of strains whose relevant genotypes/features are indicated on top and perturbations, if any, at left; strains designated Δ*Nil*, Δ*2*,*3*, or Δ1 were also Δ*infB*. Phleo supplementation was at 2 μg/ml, and I-SceI refers to Ara-supplementation in cultures of derivatives carrying P_*ara*_::I-SceI and the cognate cut site in *lacZ*. In each sub-panel, the percentage of total cells whose intensity of propidium iodide staining exceeded the threshold that was taken to demarcate dead cells (3 × 10^3^ arbitrary units, marked by vertical line), is indicated at top right. Strains employed were (all strain numbers are prefixed with GJ): I. derivatives without P_ara_::I-SceI – Δ*Nil*, 19193; Δ*2*,*3*, 19194; Δ*1*, 15494; *recA*, 19844; and II. derivatives with P_*ara*_::I-SceI – Δ*Nil*, 19804; Δ*2*,*3*, 19805; and Δ1, 19806.

### Loss of IF2-1 does not confer sensitivity to several other DSB-generating perturbations

Notwithstanding the sensitivity of the Δ*1* mutant to two-ended DSB damage, the strain was tolerant to other DSB-generating perturbations to which *recA* and *recBC* mutants are known to be sensitive. Thus, type-2 DNA topoisomerase inhibitors nalidixic acid and ciprofloxacin were tolerated to equivalent extents by the strains with differential expression of the IF2 isoforms, but conferred 10^3^-fold greater lethality upon loss of RecA or of PriB (Fig. 3A; see also Supp. Fig. S1B for sensitivity to ciprofloxacin of a *recB* but not *recO* mutant). The reversal in rank order of sensitivity between Δ*priB* and Δ*1* strains, to type-2 DNA topoisomerase inhibitors on the one hand and (high-dose) radiomimetic agents on the other, suggests that repair mechanisms following exposure to these two agent categories are distinct and different, although both are RecA- and RecBCD-mediated (1, 38).

**Figure 3:**
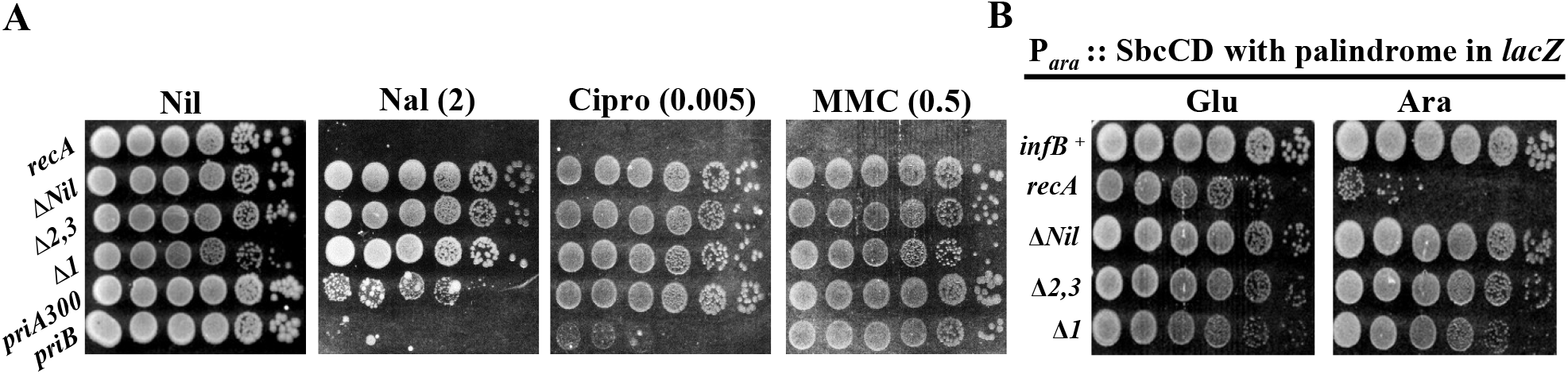
Tolerance of strains expressing different IF2 isoforms to other genotoxic agents and perturbations. Dilution-spotting assays were performed on LB with supplements as indicated on top; numbers in parentheses refer to concentrations in μg/ml. Cipro, ciprofloxacin; Nal, nalidixic acid; and MMC, mitomycin C. Relevant strain genotypes are shown at left. Strains whose designations include Δ*Nil*, Δ*1*, or Δ*2*,*3* were also Δ*infB*. In the strains of panel B, SbcCD expression that is induced in cells on Ara-supplemented medium leads to cleavage of the sister chromatid associated with the lagging-strand template at an engineered palindrome sequence in the *lacZ* gene (39). Strains used were (from top, unless otherwise indicated all strain numbers mentioned are prefixed with GJ): in panel A – 19844, 19193, 19194, 15494, 15495, and 19812; and in panel B – DL2006, 19811, 19808, 19809, and 19810.

Again unlike a *recA* derivative, the Δ*1* strain was not sensitive to mitomycin C (Fig. 3A), which cross-links DNA strands and generates DSBs following DNA replication. Leach and coworkers have described a model of site-specific DSB generation on one of the pair of sister chromatids immediately behind a replication fork, that occurs by SbcCD-mediated cleavage of a palindromic sequence (39, 40); to this perturbation as well, a strain deficient for IF2-1 remained tolerant whereas the *recA* mutant was sensitive (Fig. 3B).

### Overexpression of isoforms IF2-2,3 is also correlated with sensitivity to two-ended DSBs

In the accompanying paper (27), we have provided evidence that HR functions in *E. coli* are affected by an imbalance between the different IF2 isoforms, with the phenotypes of IF2-1 deficiency being recapitulated by overexpression in an *infB*^+^ strain of IF2-2,3 or IF2-3. With respect to DSB damage as well, our results indicate that, just as with loss of IF2-1, IPTG-induced overexpression of IF2-2,3 or of IF2-3 in the Δ*Nil* strain confers sensitivity to Phleo but not to nalidixic acid (Fig. 4).

**Figure 4:**
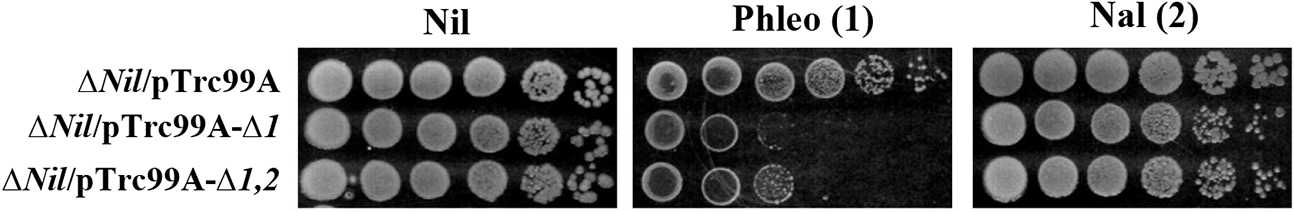
Overexpression of isoforms IF2-2,3 or IF2-3 confers sensitivity to two-ended DSBs in DNA. Dilution-spotting assays were performed on LB with Amp and IPTG, along with the additional supplements as indicated on top; numbers in parentheses refer to concentrations in μg/ ml. Nal, nalidixic acid. Relevant strain genotypes are shown at left (all were Δ*infB*). Strains used for different rows were derivatives of GJ19193 with following plasmids (from top): vector pTrc99A, pHYD5207, and pHYD5208.

### Role of IF2 isoforms in two-ended DSB repair is independent of GreA/DksA

GreA and DksA are factors modulating transcription elongation in *E. coli* (41), but they also participate in DNA repair. Loss of GreA is associated with increased tolerance to DSBs in DNA (42–44), which was confirmed for Phleo in this study (Supp. Fig. S1C, panel for 3 μg/ml). DksA is needed for DSB repair (45), and it is an apparent antagonist of GreA with respect both to DSB repair (42–44) and to other phenomena (46, 47); in our study, its loss conferred a modest sensitivity to Phleo (Supp. Fig. S1C, panels for 0.25 and 1 μg/ml).

We examined the relationship, if any, between the roles in two-ended DSB repair of GreA and DksA on the one hand, and that of IF2-1 on the other. Our results show that sensitivity to Phleo of a strain lacking IF2-1 is reversed, but only partially so, upon loss of GreA and that it is somewhat exacerbated upon loss of DksA (Supp. Fig. S1C, panel for 0.25 μg/ ml); thus the opposing effects of the two losses (IF2-1 and GreA) appear to be algebraically additive. These results suggest that the mechanism by which IF2 isoforms modulate DSB repair is different from that by GreA/DksA.

### Comparison of DNA copy number changes following site-specific two-ended DSB in Δ*Nil* and Δ*1* strains

The Herman lab (44) has previously shown by whole-genome sequencing (WGS) that following induction of synthesis of I-SceI to generate a two-ended DSB at a single genomic location in the *lac* operon, an equilibrium between DNA resection on the one hand, and re-synthesis by break induced replication (BIR) through replication restart mechanisms on the other, is reached by 30 minutes. The end result is a Chi-site modulated, asymmetric V-shaped dip in DNA copy number extending from ~100 kb ori-proximal to ~200 kb *ori*-distal of the DSB site. The *recA* mutant, on the other hand, exhibits extensive DNA copy number reduction (consequent to “reckless” RecBCD-mediated DNA degradation without any BIR), without an equilibrium being attained.

We performed similar WGS experiments to determine genome-wide DNA copy numbers in LB-grown cultures for strains carrying an I-SceI site at the *lacZ* locus, and in which the cognate enzyme (under P_*ara*_ control) was expressed in early exponential phase for one hour by addition of Ara as inducer; D-glucose (Glu) was used instead of Ara in the control uninduced cultures. The strains were Δ*infB* at the native locus and carried the ectopic Nakai *infB* constructs (ΔNil, Δ*1*, or Δ*2*,*3*); a *recA* derivative of the Δ*Nil* strain was also used.

Normalized copy number distributions from the cultures were determined as described in “Materials and Methods”, and all of them exhibited a bidirectional *oriC-to-Ter* gradient that is expected for cells in asynchronous exponential growth in rich medium (Fig. 5). Superimposed upon this gradient distribution were several distinct features of interest that are discussed below.

**Figure 5:**
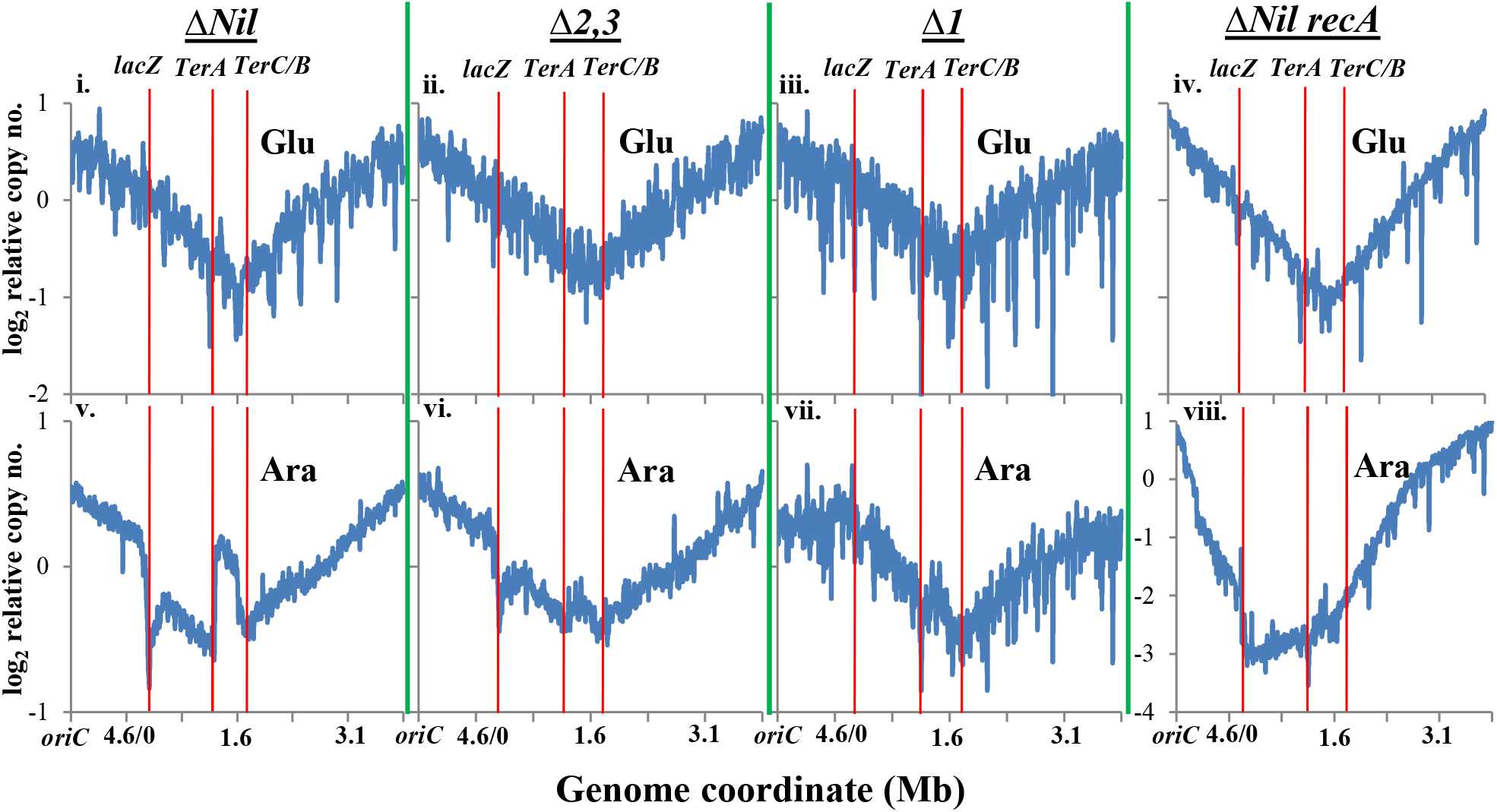
Chromosomal DNA copy number analysis by WGS in strains expressing different IF2 isoforms, following two-ended DSB generation at *lacZ*. DNA copy numbers (after normalization) are plotted as semi-log graphs for overlapping 10-kb intervals across the genome for derivatives each carrying P_*ara*_::I-SceI and the cognate cut site in *lacZ*, after supplementation of cultures grown in LB with 0.2% Glu (control) or Ara for 1 hr (top and bottom rows, respectively). Relevant genotypes are indicated at top of each pair of panels; all strains were also Δ*infB*. In these Cartesian graphical representations, the circular 4642-kb long chromosome is shown linearized at *oriC*, with genome coordinates on the abscissa corresponding to the MG1655 reference sequence (wherein *oriC* is at 3926 kb). Ordinate scales (log_2_) shown at left on top and bottom rows are common for, respectively, panels i-iv and v-vii. The positions of *lacZ*, *TerA* and *TerC/B* are marked. Strains used were (all strain numbers are prefixed with GJ): Δ*Nil*, 19804; Δ*2*,*3*, 19805; Δ*1*, 19806; and Δ*Nil recA*, 19818.

The maximum extent of copy number reduction around *lacZ* in each of the Ara-grown cultures is presented as a log_2_ dip value, in Table 1 (for derivatives of Δ*Nil* and Δ*1*). Following I-SceI induction with Ara (1-hr exposure), the Δ*Nil* strain exhibited the asymmetric V-shaped Ter-biased reduction around *lacZ* (Fig. 5 v, and Supp. Fig. S3 i; log_2_ dip = 0.8), as had previously been reported by the Herman group (44). The Δ*2*,*3* strain behaved similarly to Δ*Nil* for copy number changes around *lacZ* (Fig. 5 vi and Supp. Fig. S3 ii; log_2_ dip = 0.6). The *recA* derivative of Δ*Nil* exhibited a pattern very similar to that described by the Herman group (44), that is, a very extensive degradation on either side of *lacZ* in the former and a less drastic drop in copy number in the latter (Fig. 5 viii; log_2_ dip = 3.2). The *recA* mutant also showed a small dip in *lacZ* region read counts in the Glu-grown culture (Fig. 5 iv and Supp. Fig. S3, compare iv and v), suggestive of I-SceI cleavage in a minor proportion of cells even under uninduced conditions which is likely efficiently repaired in the Δ*Nil* strain but is lethal in *recA*.

**TABLE 1.**
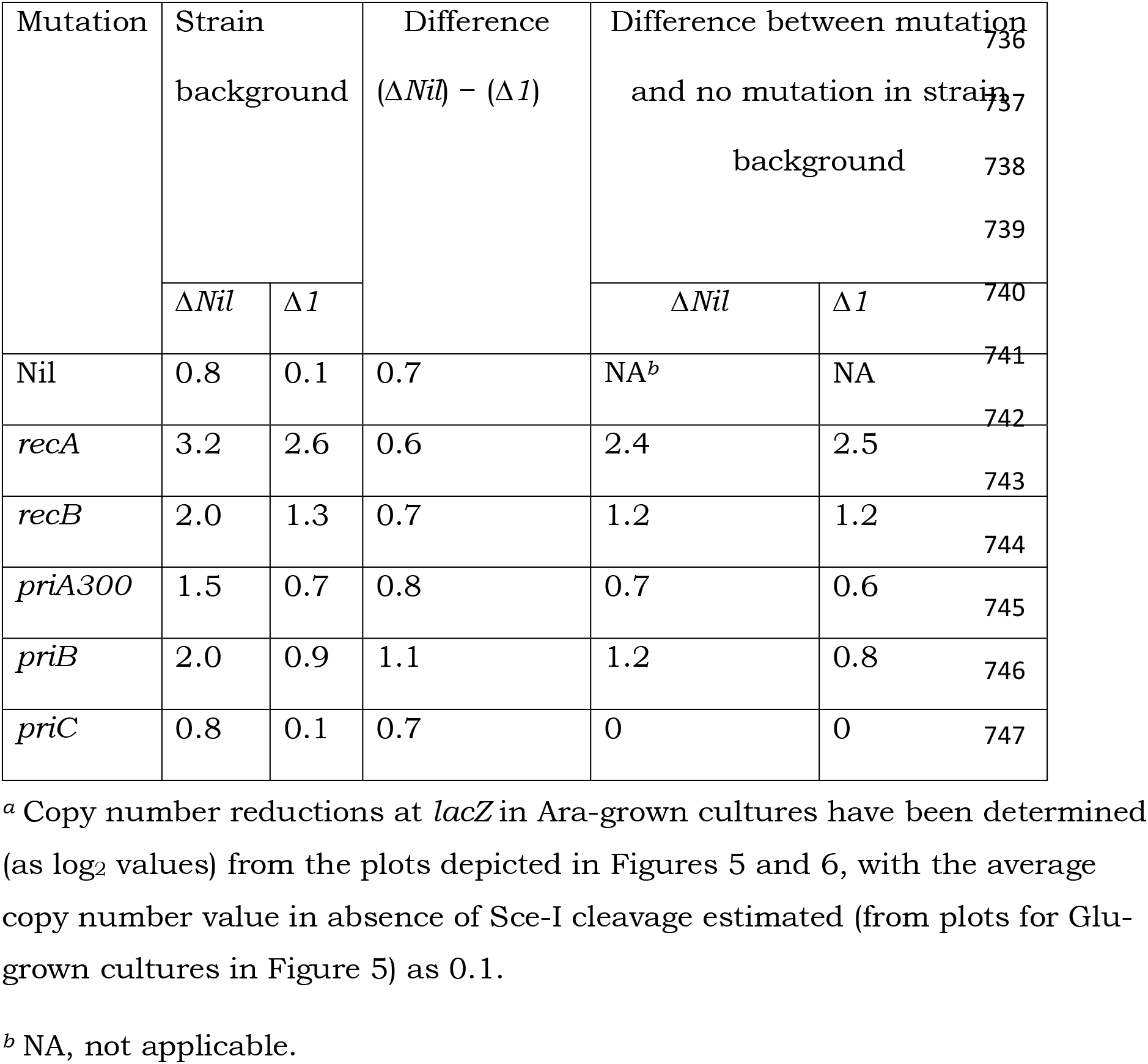
Copy number reduction (log_2_) at *lacZ* following I-SceI cleavage^a^

On the other hand, the Δ*1* strain exhibited only a very minimal reduction in read counts at the *lacZ* region (Fig. 5 vii and Supp. Fig. S3 iii; log_2_ dip = 0.1). The latter finding was quite unexpected, since it was opposite to that in *recA* with which Δ*1* shares the phenotype of pronounced sensitivity to two-ended DSBs. To exclude alternative more trivial explanations, we verified that the proportion of suppressors in the Δ*1* cultures (that is, survivors after Ara addition) was < 1%, and that the DNA sequence data for these cultures revealed no mutations in any of the candidate genes related to DNA recombination and repair.

DNA copy number determinations through WGS are considered to be quite robust (39, 48–50) in view of (i) the wealth of sequence data generated from each experiment and the simplicity of their analysis, as well as (ii) the ability to identify (from the DNA sequence) the existence of suppressor mutations if any in the cultures. As expected, therefore, the observations on copy number changes upon 1-hr Ara exposure of the Δ*Nil*, Δ*1*, or Δ*2*,*3* strains were reproducible, in that they were replicated in other independently grown cultures of these strains (Supp. Fig. S4A i-iii, respectively); furthermore, the Δ*1* strain cultured continuously with Ara also showed less or no dip in read counts around *lacZ* compared to that in the Δ*Nil* strain similarly cultured (Supp. Figs. S3 and S4B, compare, respectively, sub-panels vi with vii and i with ii).

### Contributions of DNA resection and of BIR to the copy number changes following a site-specific two-ended DSB

To dissect the relative contributions of DNA resection and of BIR to the copy number changes described above, we performed additional WGS experiments (following Ara-induced cleavage at the I-SceI site in *lacZ*) of the following single mutant derivatives of both Δ*Nil* and Δ*1* strains: Δ*recB*, *priA300*, Δ*priB*, or Δ*priC*. A Δ*recA* derivative of Δ1 was also tested, which displayed a log_2_ dip in copy number of 2.6 in the vicinity of *lacZ* (Fig. 6A i), implying DNA degradation.

**Figure 6:**
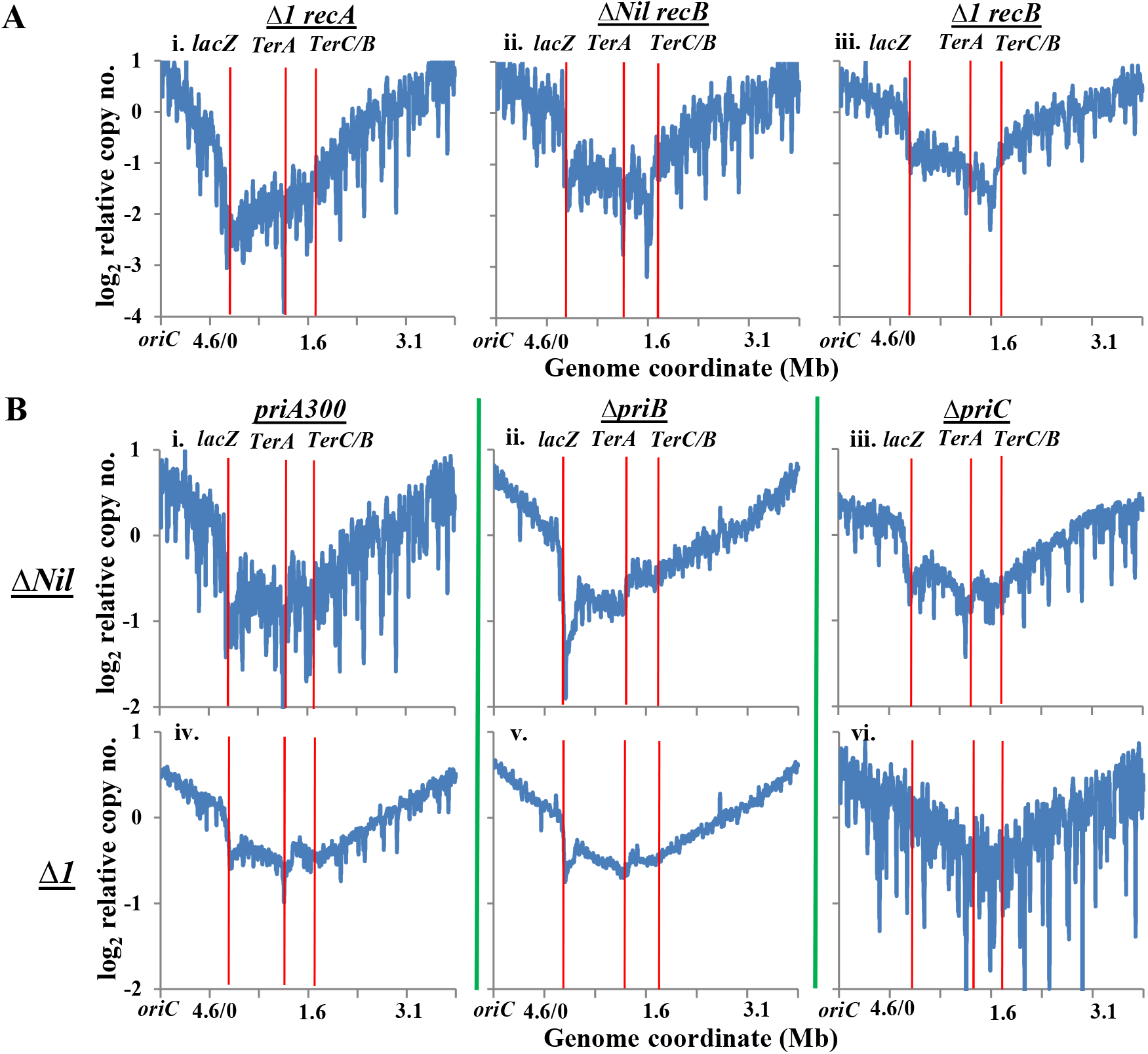
Effects of *rec* **(A)** or *pri* **(B)** mutations on DNA copy numbers in Δ*Nil* or Δ*1* strains following two-ended DSB generation at *lacZ*. Each derivative carried P_*ara*_::I-SceI and the cognate cut site in *lacZ*, and cultures grown in LB were supplemented with Ara for 1 hr. Representations of WGS analysis and notations used are as described in legend to Figure 5. Ordinate scales (log_2_) shown at left are common for all sub-panels in that row. Strains used for the different sub-panels were (all strain numbers are prefixed with GJ): (A) i, 19857; ii, 19861; and iii, 19863; and (B) i, 19864; ii, 19858; iii, 19867; iv, 19866; v, 19860; and vi, 19869.

In earlier studies from the Herman lab (44), the *recB* mutant had exhibited a substantial dip in copy number around *lacZ*; this was an unexpected finding, and was attributed to the actions (in absence of RecBCD) of RecJ and other as yet unidentified exonucleases without any BIR. In our study, the *recB* derivative of Δ*Nil* also behaved similarly, with a log_2_ dip around *lacZ* of 2.0 (Fig. 6A ii). The log_2_ dip in the *recB* derivative of Δ*1* strain was lower, at 1.3 (Fig. 6A iii). Thus, it is noteworthy that the values in both *recA* and *recB* derivatives of Δ*1* were negatively offset from those in the corresponding Δ*Nil* derivatives by about the same log_2_ magnitude (0.7) as that between Δ1 and Δ*Nil* themselves (Table 1).

With respect to each of the *pri* mutants, it was expected that the extent of additional copy number reduction if any (compared to that in the isogenic *pri*^+^ strain) would reflect the magnitude of BIR deficit conferred by the cognate *pri* mutation. The log_2_ dips in copy number in the *priA300*, *priB* and *priC* derivatives of the Δ*Nil* strain were 1.5, 2.0, and 0.8, respectively (Fig. 6B i-iii), implying a log_2_ BIR deficit (obtained by subtracting the log_2_ dip of 0.8 in *pri*^+^ Δ*Nil*) of 0.7 imposed by *priA300*, 1.2 by *priB*, and nil by *priC*.

Just as had been noted above for the *rec*^+^ *pri*^+^ as well as the *recA* and *recB* derivatives, the log_2_ dips in each of the *pri* mutant derivatives of the Δ*1* strain were shallower than those in the isogenic Δ*Nil* counterparts (Fig. 6B iv-vi, and Table 1). The values were 0.7 (*priA300*), 0.9 (*priB*), and 0.1 (*priC*),and hence the corresponding BIR deficits (after accounting for the log_2_ dip in Δ*1* of 0.1) were 0.6, 0.8, and nil, respectively.

The detailed interpretations from the WGS data near *lacZ*, with respect to a possible mechanism for mediation by IF2 isoforms of two-ended DSB repair, are provided in the “Discussion”.

### Other features of interest from the WGS data

In the chromosomal terminus region, there was a peak of read counts between the *TerA* and *TerC*/*B* boundaries that was extremely prominent for the Ara-exposed cultures of Δ*Nil* and moderately so for Δ*2*,*3* and Δ1 (Fig. 5 v-vii). As proposed earlier (51), we believe that this mid-terminus peak represents the algebraic sum of read counts of two major subpopulations, in which, respectively, clockwise and counterclockwise moving forks have traversed the terminus and are paused at Tus-bound *TerC*/*B* and *TerA*.

Another feature was the presence of sharp deep dips at several genomic positions (resembling stalactite images) even in Glu-grown cultures of several different strains. The dips were especially pronounced in the Δ*1* strain, representing log_2_ drops in normalized read counts of around 3 or more (Fig. 5 iii; and see maroon lines in Supp. Fig. S5A). Similar dips were observed in copy number curves for Glu-grown cultures of (in order of their prominence) *recA* and Δ*Nil* (Fig. 5 iv and i, respectively). The positions of these dips were identical in all three cultures, to a resolution of < 2 kb; five such representative genomic locations are depicted in Supplementary Figure S5A (green and violet lines for *recA* and Δ*Nil*, respectively).

We suggest that this feature is correlated with presence in the strains of the gene encoding I-SceI, whose basal expression is perhaps associated with nickase activity (52) at specific sequences which then leads to ss-DNA gaps at these sites [since such ss-regions are not expected to be captured in Illumina WGS protocols (44)]. Indeed, dips at several of the identical locations were observed upon re-analysis of the Herman lab data (44) for uninduced cultures of wild-type and *recA* strains carrying the I-SceI gene, with those of *recA* being the more prominent (Supp. Fig. S5B i-ii; and yellow and dark blue lines, respectively, in Supp. Fig. S5A). Interestingly, the dips were least distinct for a Glu-grown culture of the Δ*2,3* strain (Fig. 5 ii, and light blue line in Supp. Fig. S5A); this last observation serves to exclude, as a possible explanation for these dips, sequence-specific bias in generation of read numbers during WGS.

## Discussion

In this study, we report that the Δ*1* strain deficient for isoform IF2-1 is markedly sensitive to perturbations that generate two-ended DSBs in DNA (which are dependent on RecA and RecBCD for repair); a similar phenotype is elicited also upon overexpression of isoforms IF2-2,3. At the same time, the Δ*1* mutant retains tolerance to other perturbations that damage DNA and whose repair is also RecA- and RecBCD-dependent (such as exposure to type-2 DNA topoisomerase inhibitors, or DSB generation on one sister chromatid behind a replication fork). The Δ*1* mutant displays a distinctive pattern of DNA copy number changes following a site-specific two-ended DSB on the chromosome. These features are discussed below to formulate a new model to account for the action of IF2 isoforms in HR and DNA damage repair.

### Can the findings be explained by an effect of IF2 isoforms on replication restart?

Nakai and coworkers (34, 35) have previously suggested that the IF2 isoforms differentially affect the replication restart pathways, and accordingly one possible explanation for our results from WGS experiments is that in the Δ*1* mutant there is inappropriate restart and (futile) templated DNA repair synthesis at a two-ended DSB site on the chromosome. However, we believe this to be an unlikely explanation.

Thus, from amongst the replication restart mutants, the profound sensitivity to two-ended DSBs observed for a Δ*1* mutant would be mimicked only by Δ*priA* or combinations such as *priB-priC* or *priA300-priB* (13, 15). Yet, several other phenotypes of these restart-deficient mutants, including sensitivity to type-2 DNA topoisomerase inhibitors or to mitomycin C, are not observed in the IF2-1 deficient strain.

Furthermore, the Nakai group (34) has shown that, at an UV-irradiation dose which led to 80% more killing of a *priB* or *priA300* single mutant relative to *pri*^+^ (thereby establishing the need for replication restart processes for damage repair under these conditions), the Δ*1* strain was not UV^S^. Therefore, not all categories of DNA repair requiring replication restart are compromised in absence of IF2-1.

Second, we have reported in the accompanying paper (27) that the Δ*1* constructs suppress *rho-ruv* and *uvrD-ruv* lethalities, with the epistasis results indicating that IF2-1 acts downstream of the RecFORQ pre-synaptic pathway and upstream of formation of D-loops or Holliday junctions. Since replication restart occurs downstream of D-loop formation, these findings also argue against a role for IF2-1 in replication restart.

Finally, as explained below, the interpretations from WGS data of different strains following I-SceI cleavage do not also support the suggestion that loss of IF2-1 interferes with replication restart pathways.

### Interpretations from DNA copy number changes around two-ended DSB site in different strains

The Δ*Nil* (parent) strain suffered a log_2_ dip of 0.8 following 1-hr exposure to site-specific endonuclease action, which represents the equilibrium between DNA cleavage and degradation on the one hand and BIR on the other (44). The Δ*1* mutant exhibited a log_2_ dip of just 0.1, and this difference from the parent could be a consequence of decreased degradation, increased BIR, or both.

We begin with a reasonable assumption that in the *recA* and *recB* derivatives, DNA synthesis makes no contribution to copy number changes following I-SceI cleavage (since D-loops cannot occur to initiate BIR). The log_2_ dips in the *recA* or *recB* derivatives of the parent (Δ*Nil*) strain are deeper than those in the cognate derivatives of Δ*1* by the same value of around 0.7 as that seen between the *rec*^+^ pair itself. These results therefore suggest that following a two-ended DSB, the Δ*1* strain is partially protected from DNA degradation to a log_2_ extent of 0.7-fold, and furthermore that this degree of postulated protection is itself a sufficient explanation for the observed differences between the Δ*Nil* and Δ*1* strains.

Next, copy number changes around the DSB site in the different *pri* mutants can be interpreted on the assumptions that (i) the mutants are unaffected for DNA resection by RecBCD and synapsis by RecA; (ii) the pathways for BIR are redundant; and (iii) not all pathways are abolished in any of the single mutants studied. The last of these assumptions is supported by the fact that I-SceI cleavage was not lethal in single *pri* mutants of the Δ*Nil* strain. It may therefore be expected that at equilibrium, there would merely be a more pronounced dip in copy number than that in the parent, whose magnitude reflects the contribution to BIR of the cognate Pri function.

Our data suggest that BIR following a two-ended DSB in the wild-type strain is performed redundantly and additively by two pathways which are abolished, respectively, in absence of PriB and of PriA helicase. Their log_2_ contributions to copy number restoration are 1.2 and 0.7, respectively, whereas PriC makes no contribution to BIR in this situation.

Given that the net log_2_ dip in copy number in Δ*Nil* was 0.8, the gross log_2_ reduction caused by DNA resection in the strain is computed (to a first approximation) to be 2.7 (since aggregate log_2_ BIR from the two restart pathways is calculated as 1.9). This calculated value for DNA degradation is consistent with the observed log_2_ dip in *recA* of 3.2 (where there is no BIR).

Finally, all of the data on copy number changes around the DSB site in the Δ*1* derivatives may be explained on the postulates that, in absence of IF2-1, DNA degradation is decreased by a log_2_ factor of around 0.7 and that BIR capacity is more or less preserved. However, this BIR does not lead to repair of the DSB. Thus, in the Δ*1* background, the log_2_ net contributions of the PriB and PriA helicase pathways for BIR are 0.8 and 0.6, respectively, compared to the corresponding values of 1.2 and 0.7, respectively, in the Δ*Nil* parent.

### A model for the role of IF2 isoforms in two-ended DSB repair

Before formulating a model, we highlight the following observations concerning the Δ*1* strain: (i) it is as sensitive as *recA* to two-ended DSBs on the chromosome; (ii) at the same time, it is tolerant to DSBs generated by type-2 DNA topoisomerase inhibitors, mitomycin C, or palindrome cleavage on a sister chromatid behind a replication fork; and (iii) it is reasonably proficient for BIR at a two-ended DSB site, unlike the *recA* mutant.

We postulate that the fundamental defect in absence of IF2-1 is reduced annealing strength of a RecA-bound nucleoprotein filament to a target homologous DNA duplex. We further propose that even as this handicap confers a modest deficiency in all HR reactions (both RecFOR- and RecBCD-mediated) as reported in the accompanying paper (27), it has a particularly severe effect on *Ter-to-oriC* directed replisome assembly and progression (which is required only during two-ended DSB repair) (Fig. 7).

**Figure 7:**
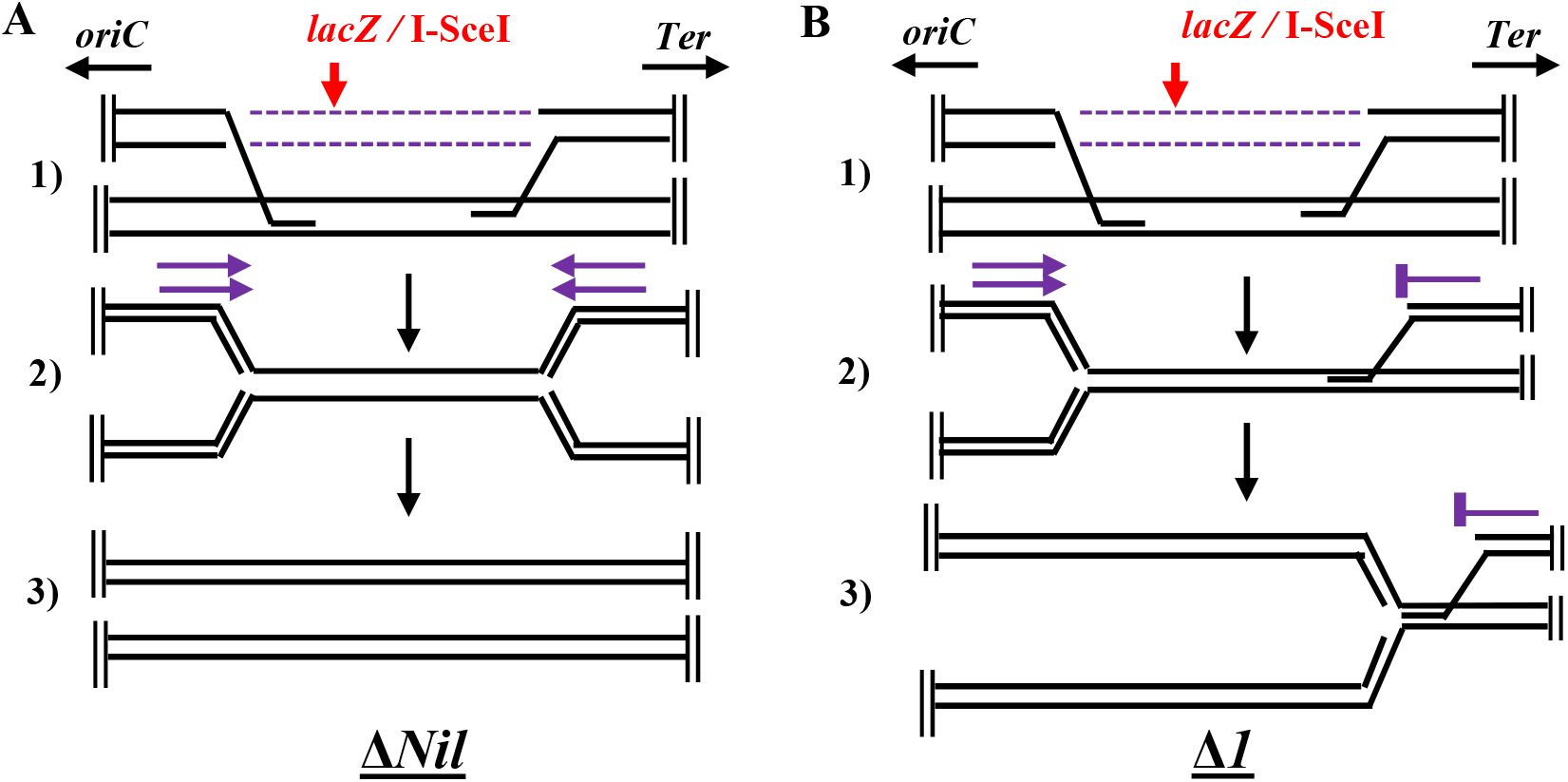
Model to explain failure of two-ended DSB repair in absence of IF2-1. **(A)** In Δ*Nil* strain, cleavage at I-SceI site in *lacZ* (red arrow) leads to DNA resection by RecBCD of the two ends towards *oriC* and *Ter*, respectively (interrupted lines), followed by two events of RecA-mediated strand invasion into a sister DNA duplex (sub-panel 1). “Ends-in” replication is initiated by redundant restart pathways (violet arrows) that are, respectively, PriA helicase- and PriB-dependent (sub-panel 2), finally resulting in successful reconstitution of two intact chromosomes (sub-panel 3). **(B)** In Δ*1* mutant, steps until initiation of RecA-mediated strand invasion are similar to those above (sub-panel 1). The *oriC*-to-*Ter* directed replisome is established through the redundant restart pathways, but that from *Ter*-to-*oriC* is not assembled (sub-panel 2). Copy number at *lacZ* is restored by the former, but two-ended DSB repair fails to be consummated (sub-panel 3).

The assumption is that a RecA nucleoprotein filament assembled on a *Ter*-proximal DSB end is more likely (in comparison with its counterpart on the *ori*-proximal DSB end) to encounter oncoming RNA polymerase molecules when it attempts to anneal into a target sister chromosome; this premise is based on the fact that a majority of heavily transcribed genes are co-directionally oriented with replication (51). In the Δ*1* mutant, such encounters would serve to majorly reduce D-loop formation. Hence, the efficiency of replisome assembly at the *Ter*-proximal DSB end will be greatly diminished; that at the *ori*-proximal DSB end would also be decreased, but to much smaller extent (note that the combined log_2_ BIR contribution in Δ*Nil* was 1.9, whereas that in Δ*1* was 1.4). Thus, although BIR will still be accomplished in the Δ*1* mutant by the *oriC*-to-*Ter* directed replisome, two-ended DSB repair remains unconsummated (Fig. 7).

Should our model be correct, the molecular mechanisms by which IF2 isoforms influence the strength of RecA-mediated annealing during HR will remain to be determined. In this context, an earlier study (53) has demonstrated that IF2 can bind DNA, through its C-terminal domain.

### Additional features in the WGS data

To explain reduced DNA degradation at a two-ended DSB site in the Δ*1* mutant, we suggest that loss of IF2-1 leads to reduced exonuclease V action [which is one component of RecBCD function (1, 5, 12)] on DSB ends. Should this postulated second effect of IF2-1 deficiency be in some way a consequence of the first (decreased annealing strength of RecA-bound DNA), it may point to existence of an interesting phenomenon of retrograde control of RecBCD nuclease function by the RecA nucleoprotein filament. A similar concept has previously been suggested for *Caulobacter crescentus* (whose RecBCD equivalent is AddAB) (54).

The other feature that distinguishes between IF2 isoforms is the set of sharp dips in copy number read counts that occur at specific genomic locations in strains with the I-SceI gene, even in uninduced cultures. On the assumption that the dips represent the net prevalence of ss-DNA gaps at these sites in cells of the population (that is, the balance between their generation and repair), it would appear that the magnitude of reduction in read counts (which is in the order Δ*1* > *recA* > Δ*Nil* > Δ*2*,*3*; Supp. Fig. S5A) inversely reflects the efficiency of their repair in the different strains. Consistent with this interpretation is our finding that loss of IF2-1 confers greater sensitivity than that of RecA to low concentrations of Phleo or Bleo (Supp. Fig. S1C), which may further suggest that in IF2-1’s absence, RecA’s non-productive binding to ss-DNA itself interferes with successful operation of the RecA-independent repair mechanisms.

### Concluding remarks

IF2 would thus represent another example, apart from NusA (55–57), DksA (43, 45), and GreA (44), of proteins earlier characterized for other critical functions also participating in DNA repair. IF2 isoforms exist in other bacteria such as, for example, in species of the Gram-negative *Salmonella, Serratia* and *Proteus* (58) as well as in Gram-positive *Bacillus subtilis* (59), and their role if any in modulating HR and two-ended DSB repair may be examined in future studies.

## Materials and Methods

### Growth media, bacterial strains and plasmids

Rich and defined growth media were, respectively, LB and 0.2% Glu-minimal A (60), and growth temperature was 37°. Concentrations used of antibiotics or Xgal were as described (27). Inducers Ara and IPTG were added at 0.2% and 0.05 mM, respectively. Genotoxic agents were added at concentrations as indicated. *E. coli* strains used are listed in Supplementary Table S1, with the following knockout alleles sourced from the collection of Baba et al. (61): *dksA*, *greA*, *recA*, *recB*, and *priC*; the Δ*infB* knockout mutation has been described earlier (62). Alleles *priA300* and Δ*priB302* were sourced from strains provided by Steve Sandler (63).

Plasmids pKD13, pKD46, and pCP20, for use in recombineering experiments and for Flp-mediated site-specific excision of FRT-flanked DNA segments, have been described by Datsenko and Wanner (64). Plasmids pHYD5207 and pHYD5208, whose construction is described in the accompanying paper (27), were used to achieve IPTG-induced expression of different IF2 isoforms from the P_*trc*_ promoter, and the parental plasmid vector pTrc99A (65) was used as control in these experiments. Plasmid pHYD5212 bearing the *infB*^+^ gene has also been described in the accompanying paper (27).

### Copy number analysis by deep sequencing after I-SceI cleavage

Strains employed each carried an I-SceI site in *lacZ* and an Ara-inducible gene construct for I-SceI enzyme (66). Following Ara-induced I-SceI cleavage, copy number determinations of the various chromosomal regions were performed by a WGS approach, essentially as described (48). After alignment of reads to the MG1655 reference sequence, the read count for each genomic region was normalized to read counts for a 600-kb region between genome co-ordinates 2501 and 3100 kb (which, relative to *oriC*, is “antipodal” to the region around *lacZ* on the opposite replichore, and is therefore expected to be the least affected following cleavage by I-SceI at *lacZ*). Additional details are given in the *Supplementary Text*.

### Other methods

Procedures were as described for P1 transduction (67) and recombineering (64). Protocols of Sambrook and Russell (68) were followed for recombinant DNA manipulations, PCR, and transformation. Procedures for flow cytometric quantitation of dead cells by propidium iodide staining (69) are described in the *Supplementary Text*.

## Supporting information

Supplementary Data including Supplementary Text, Supplementary References, Supplementary Table S1, and Supplementary Figures S1-S5.

## Data availability

The genome sequence and flow cytometry data described in this work are available for full public access from the repositories at http://www.ncbi.nlm.nih.gov/bioproject/734449 (Accession no. PRJNA734449) and https://flowrepository.org/id/FR-FCM-Z442, respectively.

## Supplemental material

Supplemental material is provided as a PDF file “Supplemental File 1”.

## Acknowledgements

We thank David Leach, Hiroshi Nakai, Susan Rosenberg, and Steve Sandler for strains; Sayantan Goswami for recombineering of the I-SceI site in *lacZ*; Nalini Raghunathan and Apuratha Pandiyan for assistance with WGS data analysis; Anjana Badrinarayanan, Rachna Chaba, Dipak Dutta, and Mohan Joshi for comments on the manuscript; and COE team members for advice and discussions.

This work was supported by Government of India funds from (i) DBT Centre of Excellence (COE) project for Microbial Biology – Phase 2, (ii) SERB project CRG/ 2018/ 000348, and (iii) DBT project BT/ PR34340/ BRB/ 10/ 1815/ 2019. JM was recipient of a DST-INSPIRE fellowship, and JG was recipient of the J C Bose fellowship and INSA Senior Scientist award.

We declare that there are no conflicts of interest.

