## Supplementary Data including Supplementary Text, Supplementary References, Supplementary Table S1, and Supplementary Figures S1-S5. for "Essential role for an isoform of *Escherichia coli* translation initiation factor IF2 in repair of two-ended DNA double-strand breaks"

### <u>Supplemental Material for</u>

Jillella Mallikarjun and J Gowrishankar#

**CONTENTS:**

**Supplementary Text.**

**Supplementary References.**

**Supp. Table S1.** List of *E. coli* strains.

**Supp. Fig. S1.** Sensitivity of strains to genotoxic agents Phleo, Bleo, and ciprofloxacin (Cipro), as determined by dilution-spotting assays.

**Supp. Fig. S2.** Effect of *recB* and *pri* mutations on tolerance to DSB generated by Ara-induced synthesis of I-SceI in strains expressing different IF2 isoforms.

**Supp. Fig. S3.** Chromosomal DNA copy number analysis by WGS in strains bearing I-SceI site in *lacZ* and expressing different IF2 isoforms.

**Supp. Fig. S4.** Chromosomal DNA copy number analysis by WGS in strains expressing different IF2 isoforms, following two-ended DSB generation at *lacZ*.

**Supp. Fig. S5.** Chromosomal DNA copy number analysis by WGS in strains carrying $P_{ara}$::I-SceI and the cognate cut site in *lacZ*, grown in absence of inducer Ara.

#### Supplementary Text

**Copy number analysis by deep sequencing after I-SceI cleavage.** All cultures were grown in LB, and two alternative protocols were adopted to achieve Ara-induced I-SceI cleavage: Ara was added to a culture in early exponential phase (at an $A_{600}$ of around 0.1) and cells were then harvested after 60 min (1), or cultures were grown to mid-exponential phase in continuous presence of Ara. Cultures grown to mid-exponential phase in LB supplemented with Glu were used as (uninduced) controls. Total DNA was extracted by the phenol-chloroform method, and paired-end deep sequencing was performed on an Illumina platform to achieve around 60- to 500-fold coverage for the different preparations. After alignment of sequence reads to the MG1655 reference sequence (Accession number NC_000913.3), gross read counts for non-overlapping 1-kb intervals were normalized to read counts per kb for the 600-kb region between genome co-ordinates 2501 and 3100 kb. The moving average method for data smoothening was as described (2).

**Flow cytometry for quantitation of dead cells in the growing cultures.** Stationary-phase cultures were subcultured at 1:5000 dilution in fresh LB medium without or with supplementation of Phleo or Ara (for induction of I-SceI) as appropriate. After growth to mid-exponential phase, 1 ml of each culture was mixed with propidium iodide at 10 μg/ml, and incubated for 15 min in the dark. Cells were then washed in 1X phosphate-buffered saline (3) before being subjected to flow cytometry. Flow cytometry was performed on the BD-FACS Aria III platform, and data (for twenty thousand cells in each preparation) were analyzed with BD-FACS Diva software (version 6.0).

**Table S1.** List of *E. coli* K-12 strains

| Strain [a] | Genotype [b] |
|---|---|
| **MG1655** | *E. coli* K-12 wild-type |
| **DL2006** | $lacI^q$ *rrnB3* Δ*lacZ4787* Δ*phoBR580* *hsdR514* Δ(*araBAD*)*567* Δ(*rhaBAD*)*508* Δ(*araFGH*) Φ(Δ$P_{araE}$ $P_{CP18}$-*araE*) Δ$P_{sbcDC}$ $P_{ara}$-*sbcDC* *lacZ*::*pal246* *cynX*::Gm |
| **GJ15410** | MG1655 Δ*lacIZYA*::FRT *galEp3 sulA11* |
| **GJ15494** | GJ15410 Δ*infB*::FRT *flgJ*::<*nusA infB*(Δ*1*)-Cm> |
| **GJ15495** | GJ15410 *priA*300 |
| **GJ15837** | MG1655 *lacZ*::<(I-SceI $_{cutsite}$)-FRT> att λ::<*araC* ($P_{ara}$-I-SceI $_{enzyme}$) –FRT> |
| **GJ19193** | GJ15410 Δ*infB*::FRT *flgJ*::<*nusA infB*(*wt*)-Cm> |
| **GJ19194** | GJ15410 Δ*infB*::FRT *flgJ*::<*nusA infB(*Δ*2,3*)-Cm> |
| **GJ19804** | GJ15837 Δ*infB*::Kan *flgJ*::<*nusA infB*(*wt*)-Cm> |
| **GJ19805** | GJ15837 Δ*infB*::Kan *flgJ*::<*nusA infB*(Δ*2,3*)-Cm> |
| **GJ19806** | GJ15837 Δ*infB*::Kan *flgJ*::<*nusA infB*(Δ*1*)-Cm> |
| **GJ19808** | DL2006 Δ*infB*::Kan *flgJ*::<*nusA infB*(*wt*)-Cm> |
| **GJ19809** | DL2006 Δ*infB*::Kan *flgJ*::<*nusA infB*(Δ*2,3*)-Cm> |
| **GJ19810** | DL2006 *flgJ*::<*nusAinfB*(Δ*1*)-Cm> |
| **GJ19811** | DL2006 Δ*recA*::Kan |
| **GJ19812** | GJ15410 Δ*priB302 cycA*::Tn*10* |
| **GJ19816** | GJ15410 Δ*recO*::Kan |
| **GJ19817** | GJ15410 Δ*recB*::Kan |
| **GJ19818** | GJ19804 *recA srl*::Tn*10* |
| **GJ19819** | GJ15410 Δ*greA*::Kan |
| **GJ19820** | GJ15410 Δ*dksA*::Kan |
| **GJ19821** | GJ15494 Δ*greA*::Kan |
| **GJ19822** | GJ15494 Δ*dksA*::Kan |
| **GJ19844** | GJ15410 Δ*recA*::Kan |
| **GJ19851** | GJ15837 Δ*infB*::FRT *flgJ*::<*nusA infB*(Δ*2,3*)-Cm> |
| **GJ19852** | GJ15837 Δ*infB*::FRT *flgJ*::<*nusA infB*(Δ*1*)-Cm> |

| | |
|---|---|
| **GJ19854** | GJ15837 Δ*infB*::FRT *flgJ*::<*nusA infB*(*wt*)-Cm> |
| **GJ19857** | GJ19852 *recA srl*::Tn*10* |
| **GJ19858** | GJ19854 Δ*priB302 cycA*::Tn*10* |
| **GJ19859** | GJ19851 Δ*priB302 cycA*::Tn*10* |
| **GJ19860** | GJ19852 Δ*priB302 cycA*::Tn*10* |
| **GJ19861** | GJ19854 Δ*recB*::Kan |
| **GJ19862** | GJ19851 Δ*recB*::Kan |
| **GJ19863** | GJ19852 Δ*recB*::Kan |
| **GJ19864** | GJ19854 *priA300* |
| **GJ19865** | GJ19851 *priA300* |
| **GJ19866** | GJ19852 *priA300* |
| **GJ19867** | GJ19854 Δ*priC*::Kan |
| **GJ19868** | GJ19851 Δ*priC*::Kan |
| **GJ19869** | GJ19852 Δ*priC*::Kan |

[a] Strains MG1655 (4) and DL2006 (5) have been described earlier; GJ15837 was from our laboratory collection constructed by Sayantan Goswami; all other strains were constructed in the present study.

[b] The following alleles and constructs have been described earlier: Δ*lacIZYA* (6); *sulA11* (7); Δ*infB*::Kan (8); att λ::<*araC* ($P_{ara}$-I-SceI $_{enzyme}$) –Cm> (9); Keio insertion-deletions (::Kan or their corresponding ::FRT derivatives) of *recA*, *recB*, *recO*, *priC, dksA*, and *greA* (10); *priA300* and Δ*priB302* (11); *cycA*::Tn*10* (12); *recA srl*::Tn*10* (13), wherein the *recA* mutation has been shown in the present study to create a Leu-to-Pro substitution at position 78 in the RecA protein; and *flgJ*::<*nusA infB*(*wt*)-Cm>, *flgJ*::<*nus AinfB*(*(*Δ*2,3*)-Cm>, and *flgJ*::<*nusA infB*(Δ*1*)-Cm>, that are designated in the text as Δ*Nil*, Δ*2,3* and Δ*1*, respectively (14).

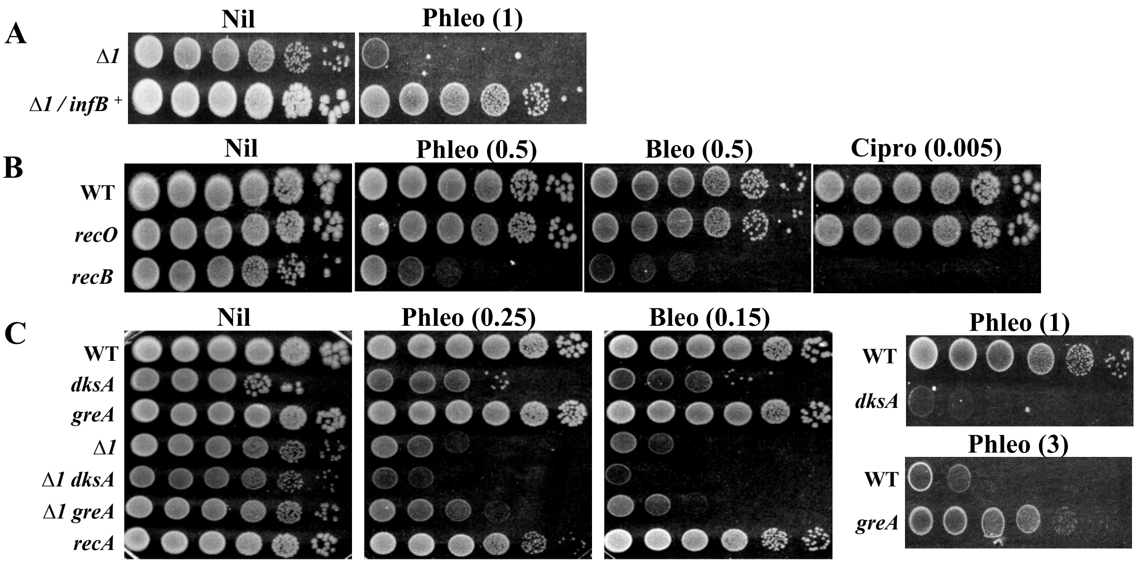


**Supplementary Figure S1**: Sensitivity of strains to genotoxic agents Phleo, Bleo, and ciprofloxacin (Cipro), as determined by dilution-spotting assays. **(A-C)** Growth medium was LB with supplements as indicated on top, and numbers in parentheses refer to concentrations in μg/ml. Relevant strain genotypes/features are shown at left; WT, wild-type. Strains whose designations include Δ*Nil*, Δ*1*, or Δ*2,3* were also Δ*infB* at the native chromosomal locus. Strains used were (all strain numbers are prefixed with GJ): WT, 15410; Δ*1*, 15494 (with plasmid pHYD5212 for the second row of panel A); *recO*, 19816; *recB*, 19817; *dksA*, 19820; *greA*, 19819; Δ*1 dksA*, 19822; Δ*1 greA*, 19821, and *recA*, 19844.

**Supp. Fig. S2**

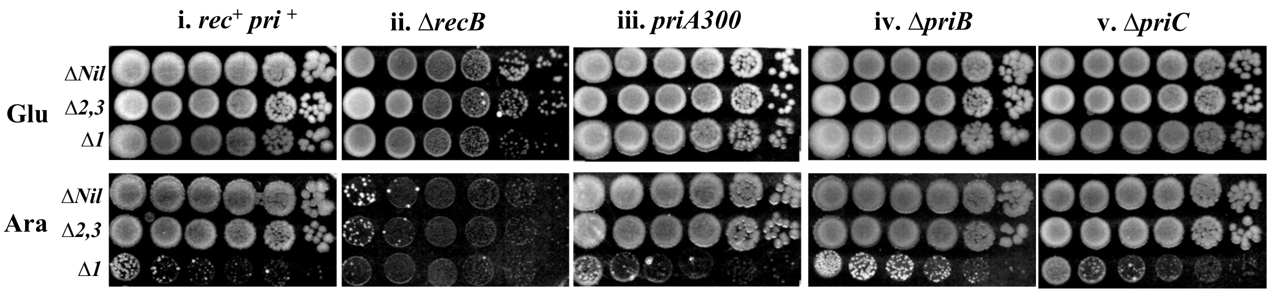


**Supplementary Figure S2**: Effect of *recB* and *pri* mutations on tolerance to DSB generated by Ara-induced synthesis of I-SceI in strains expressing different IF2 isoforms. Dilution-spotting assays were performed on LB medium supplemented with Glu (top row, control) or Ara (bottom row, test) of derivatives of Δ*Nil*, Δ*1*, or Δ*2,3* strains carrying *recB* or *pri* mutations as indicated on top of each pair of panels. All strains were also Δ*infB* at the native chromosomal locus. Note that the growth observed for *priB* and *priC* derivatives of the Δ*1* strain on Ara-supplemented medium is that of suppressor mutants. Strains used were (all strain numbers are prefixed with GJ, listed in the order *rec*$^{+}$ *pri*$^{+}$, Δ*recB, priA300*, Δ*priB*, and Δ*priC*): derivatives of Δ*Nil* – 19854, 19861, 19864, 19858, and 19867; derivatives of Δ*2,3* – 19851, 19862, 19865, 19859, and 19868; and derivatives of Δ*1* – 19852, 19863, 19866, 19860, and 19869.

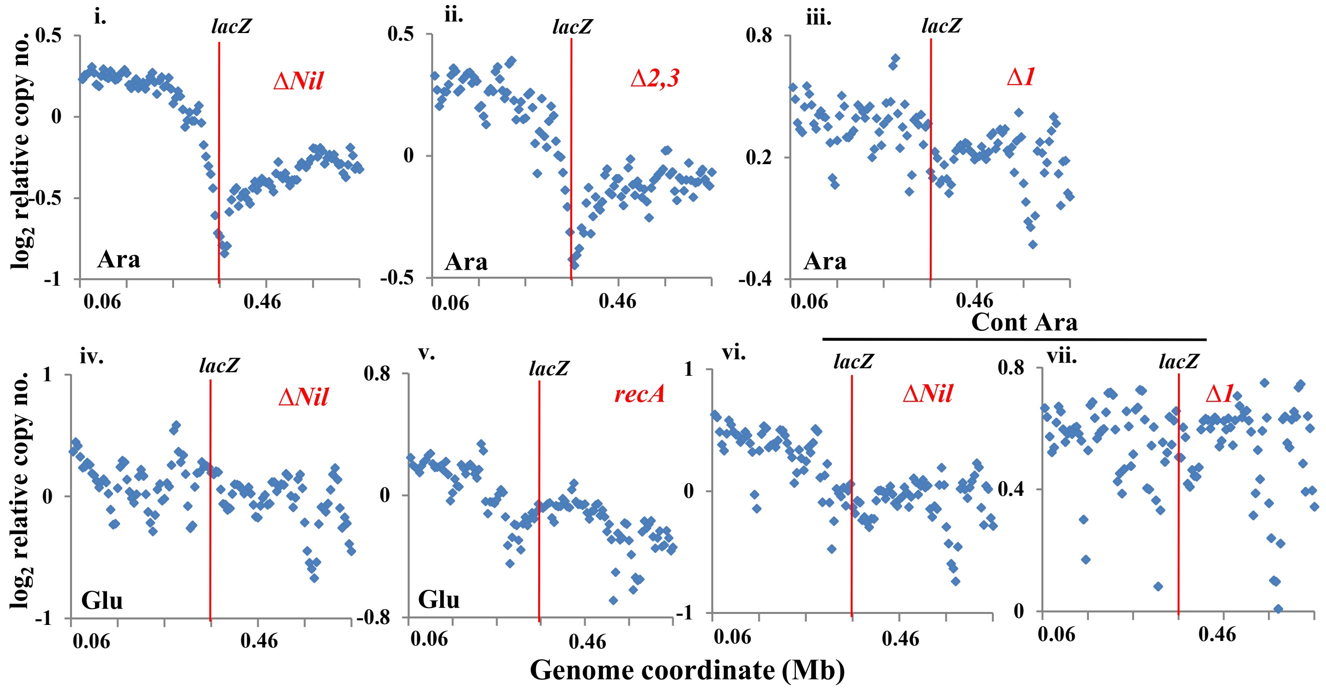


**Supplementary Figure S3**: Chromosomal DNA copy number analysis by WGS in strains bearing I-SceI site in *lacZ* and expressing different IF2 isoforms. Zoom-in images are shown for the 0.06- to 0.66-Mb genomic region (that is, 300 kb on either side of *lacZ*) in the different cultures from this study, as indicated within each of the panels. Panels i to v are from plots in sub-panels v, vi, vii, i and iv, respectively, of Figure 5; panels vi-vii are from plots in sub-panels i and ii, respectively, of Supp. Fig. S4B, for cultures grown in continuous presence of Ara (Cont Ara).

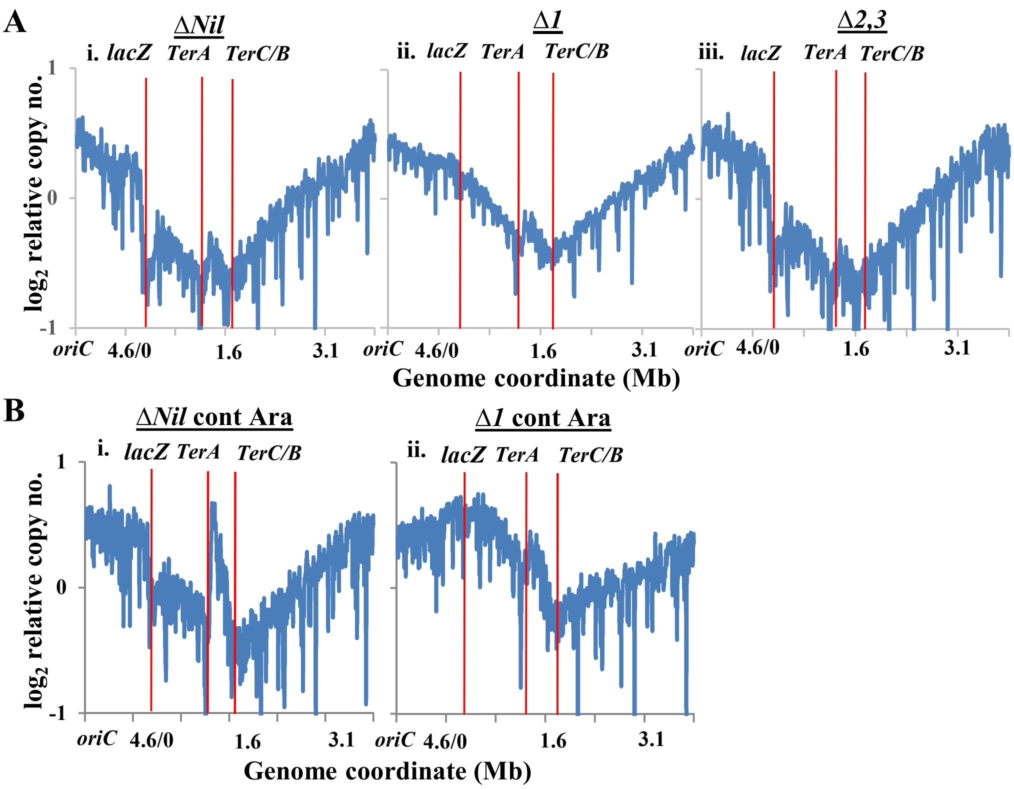


**Supplementary Figure S4**: Chromosomal DNA copy number analysis by WGS in strains expressing different IF2 isoforms, following two-ended DSB generation at *lacZ*. Each derivative carried $P_{ara}$::I-SceI and the cognate cut site in *lacZ*. Representations of WGS analysis and notations used are as described in legend to Figure 5. Ordinate scales ($log_2$) shown at left are common for all sub-panels in that row. **(A)** Plots are from cultures exposed for 1 hr to Ara, that were grown independently of those shown in Figure 5. **(B)** Plots are from cultures grown in continuous presence of Ara (Cont Ara). Strains used were (all strain numbers are prefixed with GJ): Δ*Nil*, 19804; Δ*1*, 19806; and Δ*2,3*, 19805.

**Supp. Fig. S5**

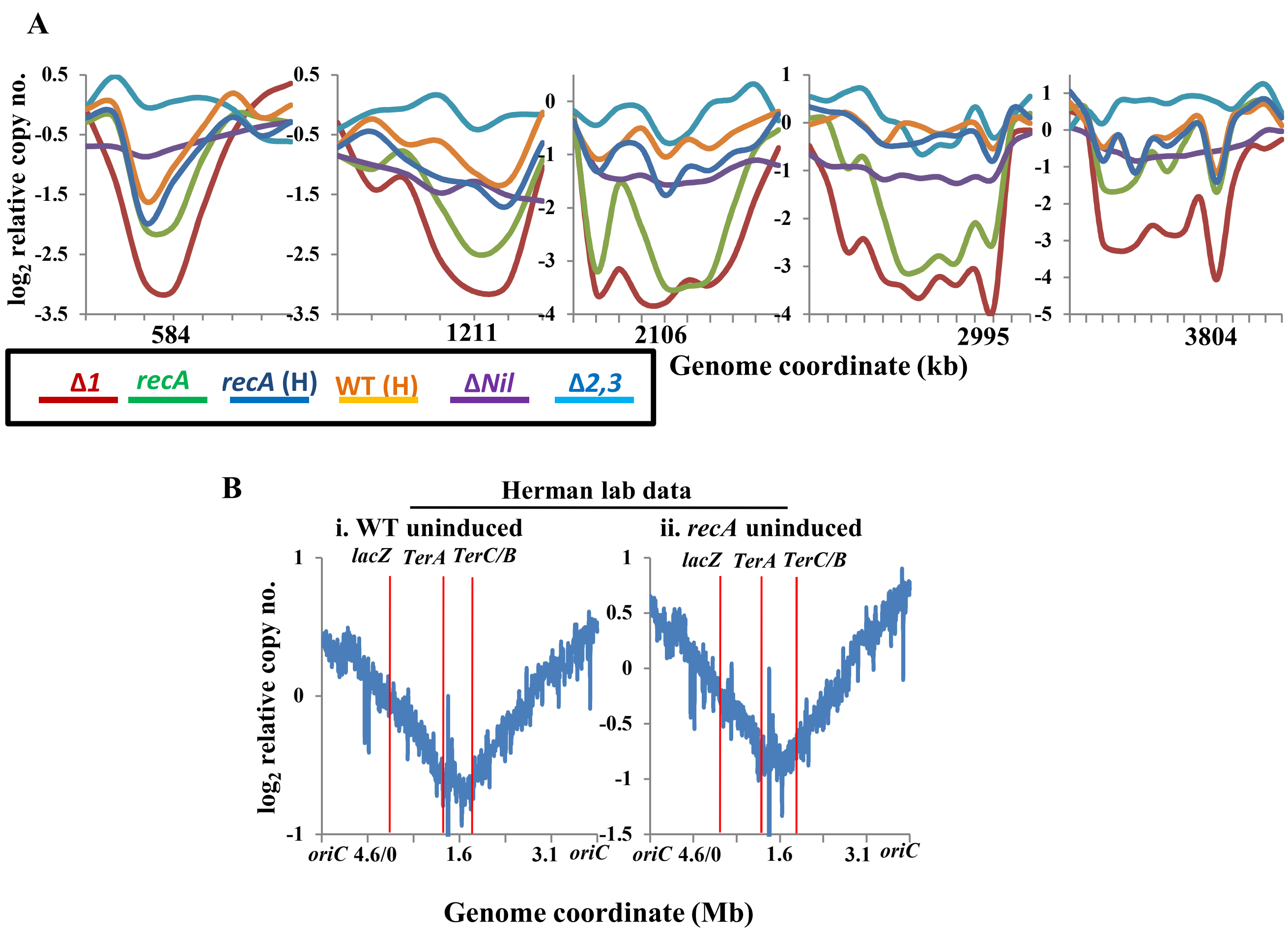


**Supplementary Figure S5**: Chromosomal DNA copy number analysis by WGS in strains carrying $P_{ara}$::I-SceI and the cognate cut site in *lacZ*, grown in absence of inducer Ara. **(A)** Zoom-in images for representative genomic regions that exhibit sharp dips in copy number. Normalized copy numbers ($log_2$) are plotted for uninduced cultures of six strains as indicated in the colours key; suffix H in parentheses identifies cultures from the Herman lab (1), whose whole-genome plot is shown in panel B below. In each sub-panel, scale markings on abscissa are 1 kb apart, and genome coordinate (in kb) corresponding to bottom of the dip is given. **(B)** Genome-wide normalized DNA copy numbers are plotted using data of the Herman lab (1) for wild-type (WT, sub-panel i) and *recA* (sub-panel ii) strains, without Ara induction. Representations of WGS analysis and notations used are as described in legend to Figure 5.
